# Enriching Biomedical Knowledge for Low-resource Language Through Translation

**DOI:** 10.1101/2022.10.11.511776

**Authors:** Long Phan, Tai Dang, Hieu Tran, Vy Phan, Lam D. Chau, Trieu H. Trinh

## Abstract

Biomedical data and benchmarks are highly valuable yet very limited in low-resource languages other than English such as Vietnamese. In this paper, we make use of a state-of-theart translation model in English-Vietnamese to translate and produce both pretrained as well as supervised data in the biomedical domains. Thanks to such large-scale translation, we introduce ViPubmedT5, a pretrained Encoder-Decoder Transformer model trained on 20 million translated abstracts from the high-quality public PubMed corpus. ViPubMedT5 demonstrates state-of-the-art results on two different biomedical benchmarks in summarization and acronym disambiguation. Further, we release ViMedNLI a new NLP task in Vietnamese translated from MedNLI using the recently public En-vi translation model and carefully refined by human experts, with evaluations of existing methods against ViPubmedT5.

## 1 Introduction

In recent years, pretrained language models (LMs) have played an important and novel role in the development of many Natural Language Processing (NLP) systems. Utilizing large pretrained models like BERT (Devlin et al., 2018), XLNET (Yang et al., 2019), ALBERT (Lan et al., 2019), RoBERTa (Liu et al., 2019), GPT-3 (Brown et al., 2020) BART (Lewis et al., 2019), and T5 (Raffel et al., 2019) has become an effective trend in natural language processing. All these large models follow the Transformer architecture proposed by (Vaswani et al., 2017) with the attention mechanism. The architecture has been proved to be very suitable for finetuning downstream tasks leveraging transfer learning with their large pretrained checkpoints. Before the emergence of large Transformer LMs, traditional wording embedding gives each word a fixed global representation. Large pretrained models can derive word vector representation from a trained large corpus. This will give the pretrained model to have a better knowledge of the generalized representation of a trained language/domain and significantly improve performance on downstream finetune tasks. The success of pretrained models on a generative domain (BERT, RoBERTa, BART, T5, etc.) has created a path in creating more specific-domain language models such as CodeBERT (Feng et al., 2020) and CoTexT (Phan et al., 2021b) for coding languages, TaBERT (Yin et al., 2020) for tabular data, BioBERT (Lee et al., 2019) and PubmedBERT (Tinn et al., 2021) for biomedical languages.

Biomedical literature is getting more popular and widely accessible to the scientific community through large databases such as Pubmed^1^, PMC^2^, and MIMIC-IV (Johnson et al., 2021). This also leads to many studies, corpora, or projects released to further advance the Biomedical Natural Language Processing field (Lee et al., 2019; Tinn et al., 2021; Phan et al., 2021a; Yuan et al., 2022). These biomedical domain models leverage transfer learning from pretrained models (Devlin et al., 2018; Clark et al., 2020; Raffel et al., 2019; Lewis et al., 2019) to achieve state-of-the-art results on multiple Biomedical NLP tasks like Named Entity Recognition (NER), Relation Extraction (RE), or document classification.

However, there have been minimum studies on leveraging large pretrained models for biomedical NLP in low-resource languages. The main reason is the lack of large biomedical pretraining data and benchmark datasets. Collecting biomedical data in low-resource languages can be very expensive due to scientific limitations and inaccessibility.

We attempt to overcome the issue of lacking biomedical text data in low-resource languages by using state-of-the-art translation works. We start with the Vietnamese language and keep everything reproducible for other low-resource languages in future work.

We introduce ViPubmedT5, a pretrained encoder-decoder transformer model trained on synthetic Vietnamese biomedical text translated with state-of-the-art English-Vietnamese translation work. Meanwhile, we also introduced ViMedNLI, a medical natural language inference task (NLI), translated from the English MedNLI (Romanov and Shivade) with human refining.

We thoroughly benchmark the performance of our ViPubmedT5 model when pretrained with synthetic translated biomedical data with ViMedNLI and other public Vietnamese Biomedical NLP tasks (Minh et al., 2022a). The results show that our model outperforms both general domain (Nguyen and Nguyen, 2020; Phan et al., 2022) and health-specific domain Vietnamese (Minh et al., 2022a) pretrained models on biomedical tasks.

In this work, we offer the following contributions:

- A first pretrained Encoder-Decoder Transformer model ViPubmedT5 that pretrained on synthetic translated biomedical data.
- A Vietnamese medical natural language inference dataset (ViMedNLI) that translated from MedNLI (Romanov and Shivade) and refined with biomedical expertise human.
- We publicize our model checkpoints, datasets, and source code for future studies on other low-resource languages.

## 2 Related Works

The development of parallel text corpora for translation and use for training MT systems has been a rapidly growing field of research. In recent years, low-resource languages have gained more attention from the industry and academia (Chen et al., 2019; Shen et al., 2021; Gu et al., 2018; Nasir and Mchechesi, 2022). Previous works include gathering more training data or training large multilingual models (Thu et al., 2016; Fan et al., 2021). Low-Language MT enhances billions of people’s daily life in numerous fields. Nonetheless, there are specific domains crucial yet limited such as biomedical and healthcare, in which MT systems have not been able to contribute adequately.

Previous works using MT systems for biomedical tasks including (Neves et al., 2016; Névéol et al., 2018). Additionally, a number of biomedical parallel (Deléger et al., 2009) have been utilized just for terminology translation only. Pioneer attempts to train MT systems using a corpus of MEDLINE titles (Wu et al., 2011), and the use of publication titles and abstracts for both ES-EN and FR-EN language pairs (Jimeno-Yepes et al., 2012). However, none of these works targets low-resource languages. A recent effort to train Vietnamese ML systems for biomedical and healthcare is Minh et al. (2022b). These, however, do not utilize the capability of MT systems, instead relying on manual crawling. Therefore, this motivation has led us to employ MT systems to contribute high-quality Vietnamese datasets that emerged from the English language. To the best of our knowledge, this is the first work utilizing state-of-the-art machine translation to translate both self-supervised and supervised learning biomedical data for pretrained models in a low-resource language setting.

## 3 English-Vietnamese Translation

Due to its limitation of high-quality parallel data available, English-Vietnamese translation is classified as a low-resource translation language (Liu et al., 2020). One of the first notable parallel datasets and En-Vi neural machine translation is ISWLT’15 (Luong and Manning, 2015) with 133K sentence pairs. A few years later, PhoMT (Doan et al., 2021) and VLSP2020 (Ha et al., 2020) released larger parallel datasets, extracted from publicly available resources for the English-Vietnamese translation.

Recently, VietAI^3^ curated the largest 4.2M highquality training pairs from various domains and as well as achieving state-of-the-art on English-Vietnamese translation^4^. The work also focuses on En-Vi translation performance across multiple domains including biomedical. The project’s NMT outperforms existing En-Vi translation models (Doan et al., 2021; Fan et al., 2020) by more than 5% in BLEU score.

## 4 Pubmed and English Biomedical NLP Researchs

The Pubmed^5^ provides access to the MEDLINE database^6^ which contains titles, abstracts, and metadata from medical literature since the 1970s. The dataset consists of more than 34 million biomedical abstracts from the literature that have been collected from sources such as life science publications, medical journals, and published online e-books. This dataset is maintained and updated yearly to include more up-to-date biomedical documents.

Pubmed Abstract has been a main dataset for almost any state-of-the-art biomedical domain-specific pretrained models (Lee et al., 2019; Yuan et al., 2022; Tinn et al., 2021; Yasunaga et al., 2022; Alrowili and Shanker, 2021; Phan et al., 2021a). Many well-known Biomedical NLP/NLU benchmark datasets are created based on the unlabeled Pubmed corpus (Doğan et al., 2014; Nye et al., 2018; Herrero-Zazo et al., 2013; Jin et al., 2019). Recently, to help accelerate research in biomedical NLP, Gu et al. (2020) releases BLURB (**B**iomedical **L**anguage **U**nderstanding & **R**easoning **B**enchmark), which consists of multiple pretrained biomedical NLP models and benchmark tasks. It is important to note that all of the top 10 models on the BLURB Leaderboard^7^ are pretrained on the Pubmed Abstract dataset.

## 5 ViPubmed

To ensure that our translated ViPubmed dataset contains up-to-date biomedical research (for example, Covid-19 diseases and Covid-19 vaccines), we use the newest Pubmed22^8^ which contains approximately 34 million English biomedical abstracts published. The raw data is compressed in XML format. We then parse these structured XMLs to obtain the abstract text with Pubmed Parser^9^(Achakulvisut et al., 2020).

By the time of our experiments, because the state-of-the-art English-Vietnamese translation NMT mentioned in Section 3 is limited to 512 token-length, we filter out abstracts with more than 512 tokens. For a fair size comparison with the unlabeled dataset of other health-related Vietnamese pretrained models (discussed in Section 8.2), we take a subset of 20M biomedical abstracts (20GB of text) for translation and leave a larger subset for future releases. We then translate the 20M English biomedical abstract with the state-of-the-art English-Vietnamese NMT released by VietAI^10^ using 4 TPUv2-8 and 4 TPUv3-8.

## 6 ViMedNLI

Along with an unlabeled dataset for pretraining, we also introduce a benchmark dataset generated by translation and refined with human experts. We start with a natural language inference (NLI) task as it less objected to errors in biomedical entity translation compared to named-entity recognition (NER) or relation extraction (RE) tasks.

### 6.1 MedNLI

MedNLI (Romanov and Shivade) is an NLI dataset annotated by doctors and grounded in the patients ‘ medical history. Given a premise sentence and a hypothesis sentence, the relation of the two sentences (entailment, contradiction, neutral) is labeled by two board-certified radiologists. The source of premise sentences in MedNLI is from MIMIC-III (Johnson et al., 2016), a large open-source clinical database. The dataset has been widely studied and benchmarked by the Biomedical NLP research community^11^ (Peng et al., 2019; Phan et al., 2021a; El Boukkouri et al., 2020; Alrowili and Shanker, 2021; Kanakarajan et al., 2019).

### 6.2 Dataset Challenges

We follow the same procedures discussed in Section 5 to translate the same training, development, and test sets released in Romanov and Shivade. The time and resources to translate the dataset are negligible as there are a total of 14522 samples.

However, upon translating the dataset with NMT, we find out that the English clinical note domain has a distinct sublanguage with unique challenges (abbreviations, inconsistent punctuation, misspellings, etc.). This observation has also been addressed in Friedman et al. (2002) and Meystre et al. (2008). Such differences in clinical language representation challenge the translation output and our quest to release a high-quality medical dataset.

### 6.3 Human Refining

The unique challenges of clinical data under translation settings (discussed in Section 6.2) require us to work with humans who not only have expertise in biomedical knowledge but are also sufficient in both English and Vietnamese languages to refine the dataset. Therefore, we collaborate with premedical Vietnamese students who studied at wellknown U.S. Universities to refine the ViMedNLI datasets.

The refining process starts with a comprehensive guidelines document with thorough annotation instructions and examples. As clinical notes contain a significant amount of technical abbreviations that the machine translation system can not translate initially (Section 6.2), we work with the medical annotators to create abbreviations and their expansion form. To make sure the expansion form of these abbreviations generalizes well in real-world settings, we verify the use case of these words through multiple Vietnamese medical websites, blogs, and online dictionaries. Hence, we decided to keep the original English abbreviations, replace them with a Vietnamese expansion form, or replace them with a Vietnamese abbreviation. Some examples of this process are shown in Figure 1.

**Figure 1:**
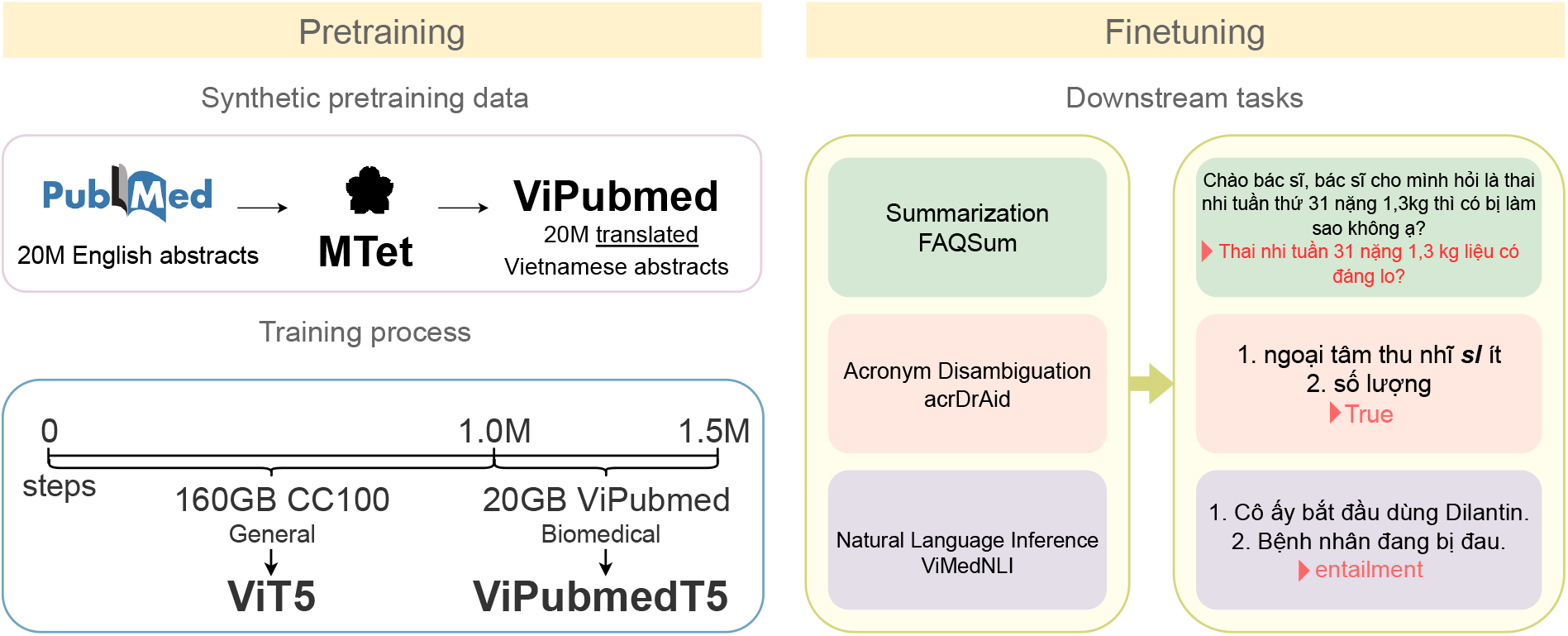
Overview of the pretraining and finetuning of ViPubmedT5

Aside from the English medical abbreviations, there are several grammatical and spelling mistakes that the machine translation system does not understand, translating either into Vietnamese meanings or even failing to translate. Human refining is therefore required. The phrase *“The infant was born at herm”*, for example, was translated as “Ðứa bé được sinh ra ở Herm”. The word *“herm”*, which should be spelled as *“term”*, is misspelled and has no medical meaning. The accurate translation should be “Ðứa bé được sinh đủ tháng”.

Additionally, the machine translation system occasionally produces incorrect Vietnamese meanings when translating words with proper English spelling and grammar. Considering the sentence *“The patient had post-term delivery”* as an example. Despite having the meaning “Bệnh nhân sinh muộn”, it was mistranslated as “Bệnh nhân sinh non” (*“The patient had pre-term delivery”*). Another example is “Narrowing of the vessels”, which actually means “Thu hẹp các mạch” rather than “Thu hẹp các”(no meaning).

## 7 ViPubmedT5

With an unlabeled synthetic translated ViPubmed Corpus (Section 5) and a benchmark ViMedNLI dataset (Section 6), we pretrain and finetune a Transformer-based language model (Vaswani et al., 2017) to verify the effectiveness of our approach in enriching Vietnamese biomedical domain with translation data. We explain our model, and the pretraining settings we applied in this section.

### 7.1 Model Architecture

We adopt the Transformer encoder-decoder model proposed by Vaswani et al. (2017), the ViT5 (Phan et al., 2022) checkpoints, and T5 framework ^12^ implemented by Raffel et al. (2019). ViT5 is the first monolingual Vietnamese Transformer model; the model achieves state-of-the-art results on multiple Vietnamese general tasks including generation and classification. The ViT5 publication releases 2 model sizes base and large. We train ViPubmedT5 using the base setting (220 million parameters) and leave larger models for future work.

### 7.2 Pretraining

We pretrain our ViPubmedT5 on 20GB of translated biomedical data ViPubmed (Section 5). We leverage the Vietnamese checkpoints in the original ViT5 work (Phan et al., 2022) and continuously pretrain the model on the synthetic biomedical-specific data for another 500k steps. Previous works (Lee et al., 2019; Tinn et al., 2021) have shown that this approach will allow pretrained language models to learn a better representation of biomedical language context while maintaining the core Vietnamese language representation.

We train ViPubmedT5 using the same spans-masking learning objective as Raffel et al. (2019). During self-supervised training, spans of biomedical text sequences are randomly masked (with sentinel tokens) and the target sequence is formed as the concatenation of the same sentinel tokens and the real masked spans/tokens.

## 8 Experiments

### 8.1 Benchmark dataset

We finetune and benchmark our pretrained ViPubmedT5 model on two public Vietnamese biomedical-domain datasets acrDrAid (Minh et al., 2022a) and our released ViMedNLI (Section 6). Detailed statistics of the three datasets are shown in Table 2.

**Table 1:** Some Examples of Abbreviations Refining

| # | MedNLI | Translated by NMT | Our Refined |
| --- | --- | --- | --- |
| 1 | Electrocardiograms revealed no QRS changes. | Điện tâm đồ cho thấy không có thay đổi về QRS . | Điện tâm đồ cho thấy không có thay đổi về phức độ QRS . |
| 2 | Patient has no PMH | Bệnh nhân không có PMH | Bệnh nhân không có tiền sử bệnh |
| 2 | Patient is post op | Bệnh nhân đã hồi phục | Bệnh nhân hậu phẫu thuật |

#1: Abbreviation in English can be used in both English and Vietnamese
#2: Abbreviation can only be used in English. In Vietnamese, abbreviation is different.
#3: Abbreviation in English is wrong when translated to Vietnamese

**Table 2:** Statistics of finetuned datasets

| Corpus | Train | Dev | Test | Task | Domain |
| --- | --- | --- | --- | --- | --- |
| acrDrAid, pairs | 4000 | 523 | 1130 | Acronym disambiguation | Medical |
| FAQSum, documents | 10621 | 1326 | 1330 | Abstractive summarization | Healthcare |
| ViMedNLI, pairs | 11232 | 1395 | 1422 | Inference | Clinical |

- **acrDrAid** (Minh et al., 2022a) is a Vietnamese Acronym Disambiguation (AD) dataset that contains radiology reports from Vinmec hospital^13^, Vietnam. The task is correctly identifying the expansion of an acronym in a given radiology report context. The dataset is annotated by three expert radiologists. The acrDrAid has 135 acronyms and 424 expansion texts in total.
- **FAQ Summarization** (Minh et al., 2022a) is a Vietnamese summarization dataset collected from FAQ sections of multiple *healthcare* trustworthy sites. For each FAQ section, the question text is the input sequence and the title is a target summary.
- **ViMedNLI** is our released dataset discussed in Section 6.

### 8.2 Baseline

In order to verify the effectiveness of our proposed methods, we compare our ViPubmedT5 model with other state-of-the-art Vietnamese pretrained models:

- **PhoBERT** (Nguyen and Nguyen, 2020) is the first public large-scale monolingual language model pretrained for Vietnamese language. The model follows the original RoBERTa (Liu et al., 2019) architecture. PhoBERT is trained on a 20GB word-level Vietnamese news corpus.
- **ViT5** (Phan et al., 2022) is the most recent state-of-the-art Vietnamese pretrained model for both generation and classification tasks. The model is trained on a general domain CC100-vi corpus.
- **ViHealthBERT** (Minh et al., 2022a) is the first domain-specific pretrained language model for Vietnamese healthcare. After initializing weights from PhoBERT, the model is trained on 25M health sentences mined from different sources.

## 9 Results

The main finetuned results are shown in Table 3. The main takeaway is that training on synthetic translated biomedical data allows ViPubmedT5 to learn a better biomedical context representation. ViPubmedT5 achieves state-of-the-art in Medical and Clinical contexts while performing slightly worse than ViT5 in healthcare topics.

**Table 3:** Tests results on Vietnamese health and biomedical tasks

| Domain | Datasets | Metrics | PhoBERT | ViT5 | ViHealthBERT | ViPubmedT5 |
| --- | --- | --- | --- | --- | --- | --- |
|  |  |  | (+news) | (+cc100) | (+health text mining) | (+translated ViPubmed) |
| Healthcare | FAQSum | R1 | 47.15 | **66.32** | 50.45 | <u>65.81</u> |
|  |  | R2 | 28.18 | **51.62** | 31.35 | <u>51.18</u> |
|  |  | RL | 41.16 | **61.3** | 43.85 | <u>60.6</u> |
| Medical | acrDrAid | Mac-F1 | 82.51 | <u>88</u> | 86.7 | **89.04** |
| Clinical | ViMedNLI | Acc | 77.29 | 77.85 | <u>79.04</u> | **81.65** |

*Notes:* The best scores are in bold, and the second best scores are underlined. PhoBERT/ViHealthBERT scores on FAQSum and acrDrAid are from Minh et al. \(2022a\)

The performance of ViPubmedT5 on the healthcare domain (FAQSum) dataset is unsurprising as the translated data from ViPubmed is biomedical scientific and has academic language representation. The FAQSum dataset, on the other hand, is mostly healthcare communication between patients and doctors on less scientific health websites.

For both medical and clinical datasets, ViPubmedT5 significantly outperforms other existing models. There are also strong improvements from the general domain ViT5 to ViPubmedT5 (88->89.04 in acrDrAid and 77.85->81.65 in ViMedNLI). This indicates that the translated ViPubmed corpus contains biomedical knowledge that low-resource Vietnamese pretrained models can leverage.

Meanwhile, our new translated ViMedNLI can serve as a strong baseline dataset for Vietnamese BioNLP research. Both health and biomedical domain models (ViHealthBERT & ViPubmedT5) perform better than general domain models (PhoBERT & ViT5) on the ViMedNLI dataset. This shows that our translated and refined ViMedNLI dataset is high-quality and has robust biomedical contexts.

## 10 Scaling to Other Languages

Our novel way of utilizing a state-of-the-art NMT system to generate synthetic translated medical data for pretrained models is not limited to the Vietnamese language and is scalable to many other low-resource languages. There are various recent works focusing on improving the quality of multiple low-resource NMT systems (NLLB Team et al., 2022; Fan et al., 2020; Bañón et al., 2020). These new state-of-the-art NMTs make the approach we discussed in this paper more practical to produce synthetic translated biomedical data, enriching the Biomedical NLP research knowledge in multiple low-resource languages.

## 11 Conclusion

We utilize the state-of-the-art translation model MTet to scale up the very low-resourced yet highly valuable biomedical data in Vietnamese. Namely, ViPubMedT5, a T5-style Encoder-Decoder Transformer pretrained on a large-scale translated corpus of the biomedical domain that demonstrated stateof-the-art results on both inference and acronym disambiguation in the biomedical domain. We also introduced ViMedNLI, a machine-translated and human expert refined benchmark in natural language inference to further grow the Vietnamese suite of benchmarks and data in biomedical data.

## 12 Limitations

Although our pretrained model trained on synthetic translated biomedical data produces state-of-theart results on downstream tasks for the Vietnamese language, the approach is hugely dependent on the quality of the NMTs for other low-resource languages. Thanks to recent studies and contributions from the Vietnamese research community (Section 3), the English-Vietnamese translation system has proven to be strong enough for us to conduct the experiments discussed in this work. However, the NMT’s actual performance needed before making the translated biomedical data useful for pretrained models is still a question that required further studies.

## Footnotes

1 https://pubmed.ncbi.nlm.nih.gov

2 https://www.ncbi.nlm.nih.gov/pmc

3 https://vietai.org

4 https://research.vietai.org/mtet

5 https://pubmed.ncbi.nlm.nih.gov

6 https://www.nlm.nih.gov/bsd/pmresources.html

7 https://microsoft.github.io/BLURB/leaderboard.html

8 https://ftp.ncbi.nlm.nih.gov/pubmed/baseline

9 https://github.com/titipata/pubmed_parser

10 https://research.vietai.org/mtet

11 https://paperswithcode.com/dataset/mednli

12 https://github.com/google-research/text-to-text-transfer-transformer

13 https://vinmec.com/

